# Preventing production escape during scale-up using an engineered glucose-inducible genetic circuit

**DOI:** 10.1101/2023.03.06.530450

**Authors:** Leonardo F. Tavares, Nathan V. Ribeiro, Vitória F. B. Zocca, Graciely G. Corrêa, Laura A. S. Amorim, Milca R. C. R. Lins, Danielle B. Pedrolli

## Abstract

A proper balance of metabolic pathways is crucial for engineering microbial strains that can efficiently produce biochemicals at an industrial scale while maintaining cell fitness. High production loads can negatively impact cell fitness and hinder industrial-scale production. To address this, fine-tuning of gene expression using engineered promoters and genetic circuits can promote control over multiple targets in pathways and reduce the burden. We took advantage of the robust carbon catabolite repression system of *Bacillus subtilis* to engineer a glucose-inducible genetic circuit that supports growth and production. By simulating cultivation scale-up under repressive conditions, we preserved the production capacity of cells, which could be fully accessed by switching to glucose in the final production step. The circuit is also resilient, enabling a quick switch in the metabolic status of the culture. Furthermore, the simulated scale-up process selected best-growing cells without compromising their production capability, leading to higher product formation at the end of the process. As a pathway-independent circuit activated by the preferred carbon source, our engineered glucose-inducible genetic circuit is broadly useful and imposes not additional cost to traditional production processes.

**GRAPHICAL ABSTRACT:** 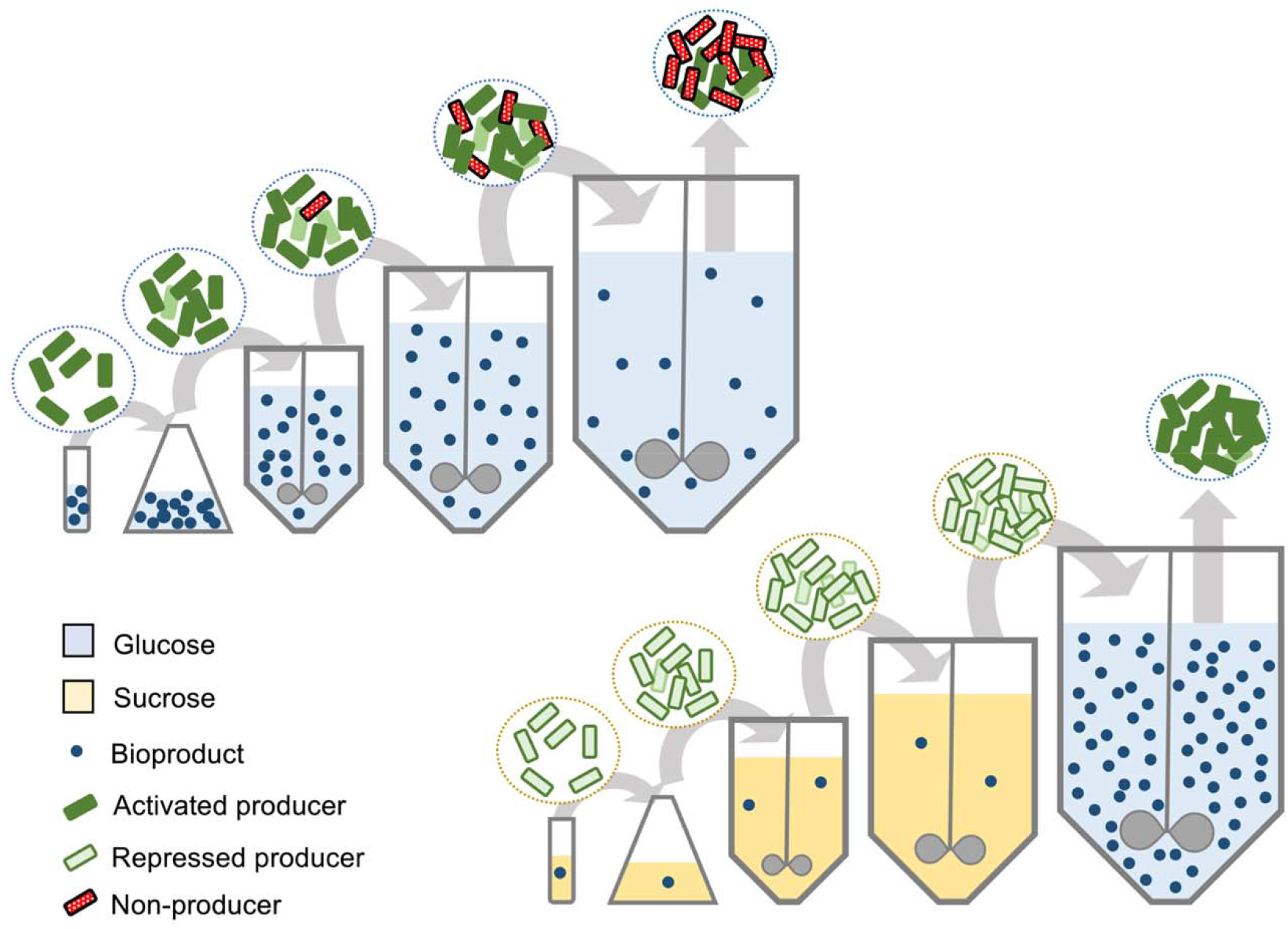

## INTRODUCTION

Synthetic biology and metabolic engineering breakthroughs in the last two decades have revolutionized the development of the bioproduction of chemicals for the benefit of industry and society. The fast development has come in times of great need as climate changes force society to rethink production ways in all sectors. The biological production of chemicals is a central contributor to the transition of the industry toward a more sustainable production chain^1^.

Large-scale industrial production of compounds from engineered microorganisms imposes some challenges such as strain instability and the emergence of non-producing cells^2^. One source of engineered cell instability comes from replicative vectors such as plasmids, which tend to be unstable during massive cell expansion to reach large-scale industrial cultivation^3,4^ Genome-engineered or edited strains cannot harbor hundreds or thousands of gene copies per cell such as multicopy plasmids can, but are more stable and preferable for large-scale production. Another source of instability is the genetic heterogeneity intrinsic to cell populations, which hinders production due to low performers^5^. Additionally, intense gene expression and target metabolite production arrest important cellular resources resulting in metabolic burden and reduced cell fitness^1^. The problem escalates if the engineered strain accumulates toxic intermediates due to different enzyme efficiencies and unbalanced pathway gene expression^4^. Compromised cell fitness is a driving force for the selection of non-producing mutant cells that naturally arise during cultivation^1,2^.

*Bacillus subtilis* is an attractive chassis for the bioproduction of chemicals for several reasons, it is a non-pathogenic and endotoxin-free bacterium, and it grows rapidly on cheap culturing medium^6^. Moreover, its potential for industrial-scale production has been long known from the riboflavin production bioprocess^7^. Finally, its high genetic stability, natural competence, and endogenous DNA recombination machinery facilitate and simplify the use of CRISPR for gene edition^8^.

Promoters play an essential role in regulating gene expression and are frequently the primary choice of control in engineered microbial strains^3,9^ Promoter engineering in *B. subtilis* has focused on developing libraries of constitutive promoters with a broad range of strengths^10–12^. Additionally, recent efforts have been directed at engineering quorum-sensing responsive promoters^13–16^. However, only a few inducer-responsive promoters are available for *B. subtilis* responding to cumate, bacitracin, IPTG, pyruvate, xylose, and maltose^17–21^. Among them, the last four induction systems are susceptible to catabolite repression by glucose, preventing the use of this preferred carbon source in the production medium. Therefore, we decided to take advantage of the catabolite repression regulatory process to create a glucose-induced regulatory circuit for industrial-scale bioproduction.

Carbon catabolic repression (CCR) is widespread in free-living heterotrophic bacteria, helping them to make the most efficient use of the carbon sources available. In *B. subtilis*, CCR regulation is mediated by the repressor protein complex CcpA-HPr(Ser-P). Feeding with preferred carbon source, such as glucose, results in high intracellular concentrations of fructose-1,6-bisphosphate and ATP, leading to HPr phosphorylation at the Ser46. The active phosphorylated complex CcpA-HPr(Ser-P) binds to palindromic operators in promoter regions called catabolic responsive elements (*cre*) preventing RNA polymerase from binding to the promoter or causing a roadblock to RNA synthesis^22,23^. We explored the well-established capacity of CCR to regulate gene expression in response to carbon feed to engineer a regulatory gene circuit to induce gene expression upon culture feeding with glucose, the preferred carbon source of *B. subtilis* and many other bacteria. As a global regulatory process, CCR can efficiently sense the availability of glucose and regulate the expression of several genes^22,24^, making it a robust tool for metabolic engineering.

## RESULTS

### Promoter engineering

We engineered the widely used *veg* promoter (P_*veg*_) to convert it from a constitutive to two regulated promoters. To engineer a carbon-responsive promoter, we added two catabolite-responsive elements (*cre* sites) downstream to the P_*veg*_ to create P_*veg-cre*_. Additionally, we added the *tetO* site downstream to the P_*veg*_ to create the TetR-responsive promoter P_*veg-tetO*_ (Figure 1A and Supplementary table 1). We measured growth and luminescence emission generated by the reporter *luxABCDE* gene cluster under the control of P_*veg*_ and its derivatives during cultivation in different carbon sources (Figure 1B-D). As expected, glucose strongly represses the gene expression driven by the P_*veg-cre*_, while lactose, xylose, and maltose activate it. Although glycerol supported growth better than lactose, it negatively impacted luminescence production driven by all three promoters (Figure 1B-D). Important to notice, the *cre* sites strongly reduced the P_*veg*_ activity even when induced by glucose. In the assay, the P_*veg-tetO*_ was tested in a strain lacking TetR and, therefore, should behave as constitutive. That was true for most carbon sources tested. However, feeding the strain with sucrose resulted in high production of luminescence (Figure 1D).

**Figure 1.**
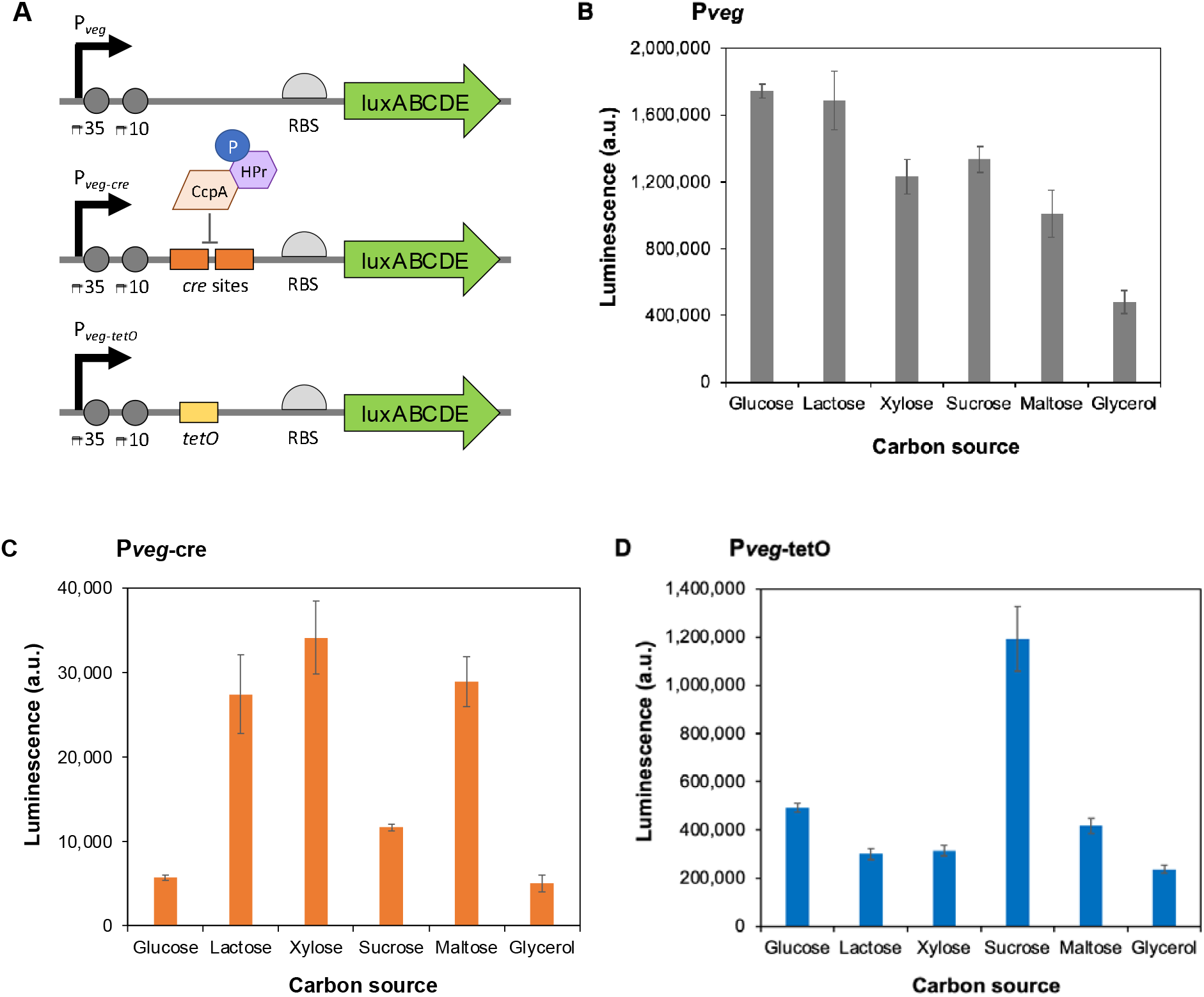
Promoter design and characterization. (A) Three different promoter designs were used to control the *luxABCDE* reporter gene cluster, the strong constitutive P_*veg*_, and the engineered promoters P_*veg-cre*_ and P_*veg-tetO*_. (B) Luminescence production under the control of t**he** P_*veg*_ by *B. subtilis* fed with different carbon sources. (C) Luminescence production under t**he** control of the P_*veg-cre*_ by *B. subtilis* fed with different carbon sources. (D) Luminescen**ce** production under the control of the P_*veg-tetO*_ by *B. subtilis* fed with different carbon sources. The strain in (D) lacks *tetR* and, therefore, the P_*veg-tetO*_ response can only result from the general metabolic condition in the cell in each condition. Results are presented as the mean of biological triplicates and the standard error. All raw data files are available in Supplementary data 1.

### Engineering a glucose-responsive circuit

The new promoters were used to engineer a glucose-responsive circuit to control gene expression. The circuit was designed to invert the signal of glucose-induced catabolic repression to create a glucose induction process for target gene expression. Therefore, *tetR* was cloned under the control of the P_*veg-cre*_, the T7 RNA polymerase gene was cloned under the control of the P_*veg-tetO*_, and the T7 promoter was cloned upstream of the *luxABCDE* gene cluster (Figure 2A). The complete gene circuit was integrated into the *B. subtilis* genome as a single copy. The resulting strain was cultivated in different carbon sources for growth and luminescence measurements (Figure 2B). As expected, glucose induced gene expression, while xylose, maltose, glycerol, sucrose, and lactose repressed it. A leakage control strain, carrying only the *luxABCDE* under the control of the T7 promoter, showed only background luminescence similar to *B. subtilis* wild-type when fed with glucose (data not shown); therefore, the luminescence measured can only be assigned to the circuit’s function. The strongest repression effects were measured for glycerol, sucrose, and lactose. The glycerol effect may result from a generally reduced promoter activity affecting the circuit, as observed for the individually characterized promoters (Figure 1B-C). Nevertheless, it supported growth and repressed gene expression properly, constituting a useful carbon source to turn off the circuit. Although lactose strongly reduced the luminescence production, the strain suffered to grow fed by it in the conditions tested. From all non-preferential carbon sources tested, sucrose resulted in the strongest repression (5.2-fold) and supported robust growth (Figures 2B-C), constituting the best source for repression of gene expression.

**Figure 2.**
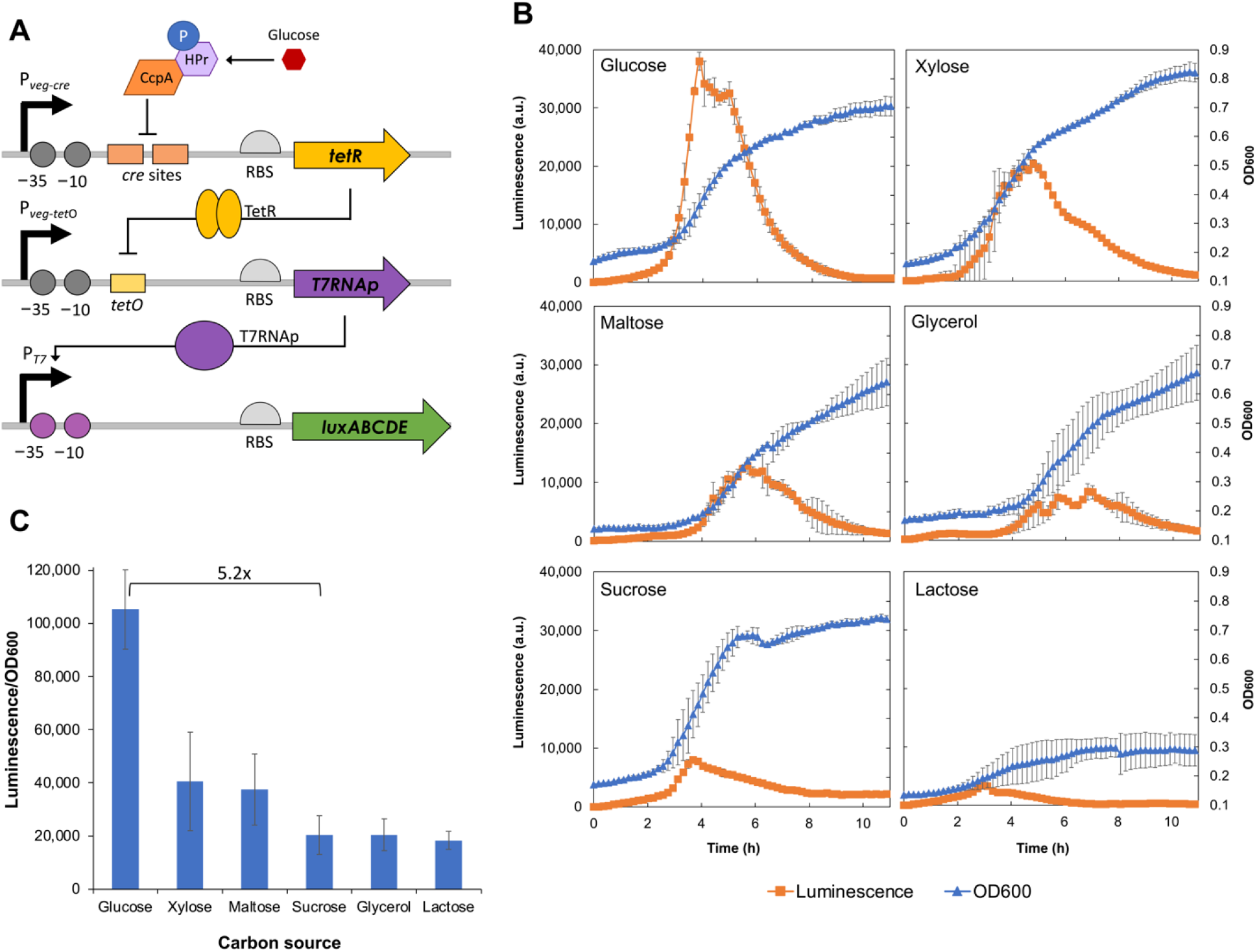
Glucose-inducible genetic circuit. (A) Circuit design. When *B. subtilis* is fed with its preferred carbon source, such as glucose, the CcpA-HPr(Ser-P) complex binds to the *cre* sites in the P_*veg-cre*_ and represses the *tetR* expression. In the absence of TetR, the T7 RNA polymerase is produced and drives the transcription of the reporter gene cluster *luxABCDE*. (B) Growth and luminescence production profile of *B. subtilis* expressing the *luxABCDE* under the control of the glucose-inducible genetic circuit fed with different carbon sources: glucose, xylose, maltose, glycerol, sucrose, and lactose. (C) Maximal specific production (luminescence/OD_600_) for each carbon source. Results are presented as the mean of biological triplicates and the standard error. All raw data files are available in Supplementary data 2.

### The glucose-induced circuit is resilient

One important feature of regulatory processes is the ability to quickly switch from one state to the other. After demonstrating the ability of our circuit to activate and deactivate gene expression in response to the carbon source available, we next tested its response to sequential cultivation in a preferred carbon source after cultivation in a non-preferred, and vice-versa. We cultivated the strain overnight in glucose or xylose and then used the cultures to inoculate new ones switching the carbon sources. The circuit quickly switched from repressed to induced (and vice versa), resulting in the expected level of gene expression for the second source (Figure 3A) and demonstrating its resilience.

**Figure 3.**
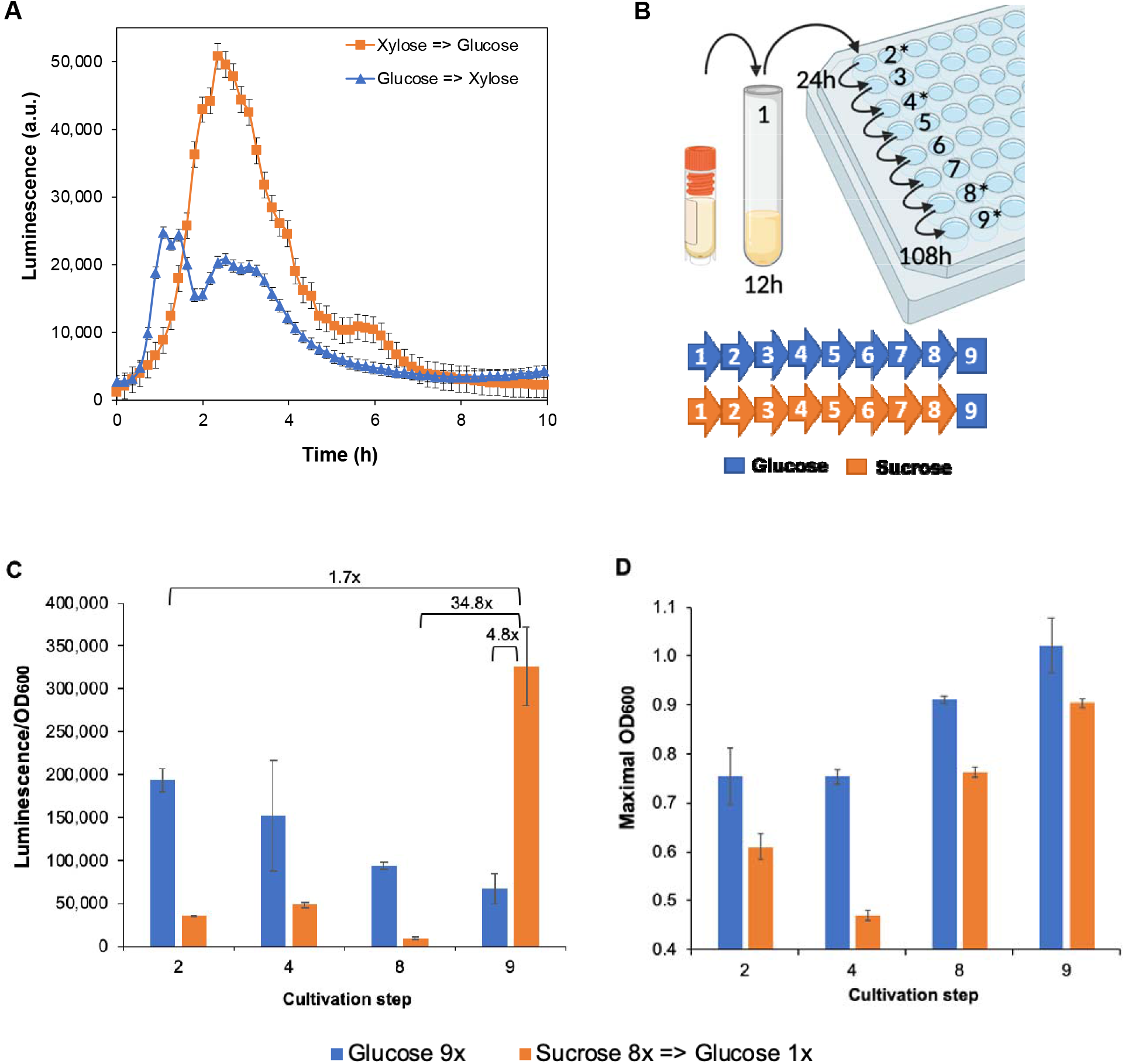
Circuit resilience and scale-up simulation. (A) Luminescence production profile under control of the glucose-induced genetic circuit submitted to carbon source change. Time zero marks the change to the second carbon source. (B) Production scale-up was simulated through nine serial cultivation transfers (≥ 55 generations) starting from glycerol stocks to a test tube followed by cultivation in a 96-well microplate. Two treatments were applied, continuous glucose feeding for nine steps and eight steps of sucrose feeding followed by a glucose feeding ninth step. The ninth step simulates the final large-scale production cultivation. Transfers were carried out every 12h, and the asterisks mark the steps with results presented in C and D. (C) Maximal specific production (luminescence/OD_600_) measured during steps 2, 4, 8, and 9 of the simulated scale-up. The strain was either fed with glucose exclusively for all nine steps or sucrose for eight steps and glucose in the ninth step. (D) Maximal OD_600_ reached by the strain in the steps and for the same treatments as in C. Results are presented as the mean of biological triplicates and the standard error. All raw data files are available in Supplementary data 3 and 4.

### Large-scale industrial production simulation

The gradual scale-up from a master stock culture to the production bioreactor in the industry can lead to a severe loss of production due to selective enrichment of nonproducing cells^1^. To further test the circuit’s resilience and to simulate the circuit behavior over industrially relevant timescales, we serially transferred the production strain every 12h for a total of nine times, accounting for at least 55 generations, and measured growth and luminescence emission (Figure 3B). The strain was fed with glucose during all cultivations for continuous production induction in one treatment. In another treatment, the strain was fed with sucrose for the first eight cultivations and switched to glucose at the last cultivation step to lower the production load during the scale-up and only fully activate gene expression in the simulated production step. The cultivation fed exclusively with glucose produced the expected luminescence after two steps (Figure 3C), which is equivalent to the cultivation performed before (Figure 2B). However, in the simulated production cultivation (step 9) there was a loss of 60% in the maximal luminescence emission. During the process, the culture’s maximal density increased by 35% (Figure 3D and Supplementary figure 1A). On the other hand, sucrose feeding repressed the production in the first eight steps and efficiently prevented production escape, leading to a 34.8-fold induction at the last step when the strain was fed with glucose (Figure 3C). Moreover, in the simulated production scale, the sucrose/glucose-fed strain generated a 4.8-fold higher luminescence/OD_600_ than the exclusive glucose-fed one, which represents the highest production measured for the strain and a 1.7-fold increase over standard 2-step cultivation in glucose. The procedure also selected the best-growing cells resulting in an increase in the culture’s maximal density of 48% (Figure 3D and Supplementary figure 1B). Noteworthy, a nonexpected drop of 25% in the culture maximal density was observed during the fourth step in the sucrose-fed cultures. Yet in the next steps, the cultures recovered and increased their growth capacity.

## DISCUSSION

Engineering highly productive, robust, and stable strains is still a huge challenge. Success in lab-scale production may not be transferable to large-scale production. Cell metabolism has been shaped by evolution to support growth and maintenance. Therefore, engineered strains strive to keep homeostasis and counteract engineering efforts if it compromises intensive cell resources. This is especially important for strains aimed at industrial bioprocesses, which require engineered cells to grow robustly in sequential vessels of increasing volume to reach the final large-volume production fermentation^1–3^. Engineered biosynthetic pathways are commonly designed with gene expression under the control of either constitutive or induced promoters. Chemically induced promoters are widely used and are very useful during the first engineering steps for implementing a new pathway. However, the use of expensive inducers is cost-prohibitive for most industrial-scale processes. Therefore, constitutive promoters have been preferred for engineering such industrial strains^3,25^. Constitutive production often results in high yields for lab-scale production. However, it may also result in a high production load due to intense continuous gene expression and accumulation of pathway intermediates, compromising cell fitness^1,3^. Phase-dependent promoters, quorum-sensing-based control, and other dynamic tools to control gene expression are the available alternatives to reducing the metabolic burden in engineered cells and avoiding additional costs in large-scale bioprocesses^3,9,26^. Although useful, these solutions may not always meet large-scale production requirements. Ultra-deep sequencing was recently used to identify multiple recurring intra-pathway mobile element insertions (IS) in mevalonic acid-producing *Escherichia coli*. The insertions accumulated in the cell population over nine serial cultivation transfers simulating a large-scale fermentation process, resulting in an increase in the growth rate while the production was completely lost^1^. Based on their results, we designed a similar scale-up experiment to test the stability of our engineered *B. subtilis*. As for *E. coli*, *B. subtilis* serial cultivation resulted in production cells being selected against, in competition with more fit non-producing cells when production was continuously induced by glucose. By lowering the production load during the scale-up process, we were able to not only prevent loss in the production capacity but also increase it. The serial transfer cultivations in sucrose selected the best-growing cells increasing the maximal culture density as seen in glucose. Possibly, random mutations in the production pathway during sucrose cultivation did not result in any substantial gain in fitness, as the production load was low. Therefore, the growth increase in the last step might have been caused by the selection of mutations in production non-related loci. As the fittest cells selected in sucrose were still fully capable of production, the final cultivation in glucose delivered the highest production level measured for the strain.

Genetic heterogeneity of engineered cells and loss of production during serial cultivations have been addressed in *E. coli* and *S. cerevisiae* before. *E. coli* was reported to lose its mevalonic acid production after 95 generations^27^, and *S. cerevisiae* lost most of its vanillin-β-glucoside production after only 25 generations^28^ when no control was applied. Production loss could be reverted using a pathway-specific biosensor to couple product formation to the expression of essential genes^27,28^ or to the expression of an antibiotic resistance gene combined with continued use of the antibiotic in the culture^5^. The use of fatty acid and tyrosine sensor-regulators to control the *tetA* expression resulted in the selection of the best *E. coli* producers for each end-product under tetracycline pressure, increasing production up to 2.6-fold^5^. For long-term performance, a mevalonic-acid-responsive regulator coupled with the expression of essential genes extended the productive lifetime of *E. coli* for 95 generations without any change in growth rate^27^. A similar approach was taken for *S. cerevisiae* producing vanillin-β-glucoside, but using sensors for pathway intermediates. The resulting strain remained productive for at least 55 generations^28^. Although efficient, these sensor-regulator circuits are limited to specific metabolites, and in one case there is a need for antibiotics in the culture. On the other hand, our regulatory circuit is not pathway-specific and can be widely applicable to the production of any molecule in *B. subtilis*. Moreover, the concept may be expanded to other Firmicutes that use a similar CCR machinery^22^.

## CONCLUSION

We engineered a glucose-repressed promoter from the strong P_*veg*_ and *cre* sites from *B. subtilis*. The resulting promoter, P_*veg-cre*_, was further used to engineer a glucose-induced genetic circuit that is useful for metabolic engineering of industrial bacterial strains. The circuit is resilient and can be used to reduce the production load during scale-up to large bioreactor volume; therefore, preventing undesired production scape. Moreover, the circuit is broadly useful for the production of any molecule in *B. subtilis* and potentially in other bacteria. We also envision utility for our circuit in engineered living therapeutics^29,30^ and biosensors^31^.

## MATERIALS AND METHODS

### Plasmids and other DNA sequences

The plasmids used were derived from the *Bacillus BioBrick* Box: pBS3C*lux* (BBa_K823025) and pBS4S (BBa_K823022)^17^. Cloning work has been performed using the standard BioBrick assembly method. *tetR* and T7 RNA polymerase genes were PCR amplified using as templates the plasmids pSB1C3-TetR-LVA (BBa_P0440) and pSB1C3-T7RNAP (BBa_K145001), respectively. Both PCR products were ligated into the pBS4S, generating two separate plasmids. Promoter sequences from P_T7_, P_*veg*_, P_*veg-cre*_, and P_*veg-tetO*_ were constructed from oligonucleotides forward and reverse strands designed to form overhangs when annealed that were compatible with the BioBrick assembly. The PT7 and P*veg* were ligated into the pBS3Clux separately, generating the plasmids pBS3C-PT7-lux and pBS3C-P_*veg*_-lux. P_*veg-cre*_ was ligated into the pBS4S-tetR-LVA, which coded for a TetR fused to the LVA degradation tag. P_*veg-tetO*_ was ligated into the pBS4S-T7RNAP to generate pBS4S-P_*veg-tetO*_-T7RNAP. The P_*veg-tetO*_-T7RNAP was the PCR amplified and ligated into the pBS4S-P_*veg-cre*_-tetR-LVA to generate the pBS4S-P_*veg-cre*_-tetR-LVA-P_*veg-tetO*_-T7RNAP. All constructions were confirmed by Sanger sequencing. All the plasmids used in this work are listed in Supplementary table 2.

### Bacterial strains, transformation, and growth conditions

*E. coli* Top10 was used for cloning and propagation of plasmids. It was aerobically cultivated at 37°C in Lysogeny Broth (LB) enriched with 100 μg mL^-1^ ampicillin when needed. *E. coli* was transformed with the plasmids using a standard heat-shock protocol and in-house produced chemically competent cells. *B. subtilis* KO7 (BGSCID 1A1133), a protease-free host, was used as chassis for the experiments and was routinely cultivated in LB medium for propagation. *B. subtilis* was transformed with the plasmids using the Two-step *Bacillus subtilis* Transformation Procedure. After transformation, sequences were PCR amplified from genomic DNA for chromosomal integration confirmation and submitted to Sanger sequencing. For growth and luminescence measurements, *B. subtilis* strains were cultivated aerobically in the PW complex medium: 1% carbon source (m/v for solids and v/v for glycerol), 1 g L^-1^ yeast extract, 25 g L^-1^ NaNO_3_, 0.333 g L^-1^, KH_2_PO_4_, 1 g L^-1^ Na_2_HPO_4_·12H_2_O, 0.15 g L^-1^ MgSO_4_·7H_2_O, 7.5 mg L^-1^ CaCl_2_, 6 mg L^-1^ MnSO_4_·H_2_O, 6 mg L^-1^, FeSO_4_·7H_2_O, pH 7.0). When required, 5 μg mL^-1^ chloramphenicol and/or 100 μg mL^-1^ spectinomycin was added to the culture medium. All the strains used in this work are listed in Supplementary table 3.

### Growth and luminescence measurements

All measurements were performed on a microplate reader Tecan Infinite 200 Pro (Tecan, Männedorf, Switzerland) with a working volume of 200 μL in 96-well plates with optically clear bottoms. Cell growth was monitored by optical density at 600 nm wavelength (OD_600_). Luminescence was measured on white plates using an integration time of 1,000 ms. The microplate reader was set at 37°C and orbital shaking (143 rpm), and measurements were taken every 10 min. All experiments were carried out using biological triplicates.

## Author Information

Danielle B. Pedrolli: Universidade Estadual Paulista (UNESP), School of Pharmaceutical Sciences, Department of Bioprocess Engineering and Biotechnology, Rodovia Araraquara-Jau km1, 14800-903 Araraquara, Brazil Milca R.C.R. Lins: Federal University of ABC (UFABC), Center for Natural and Human Sciences, Campus Santo Andre, Avenida dos Estados 5001, 09210-580 Santo André, Brazil

## Author Contribution

L.F.T.: investigation, validation, data curation, and analysis. N.V.R.: conceptualization and investigation. V.F.B.Z.: analysis and writing - review & editing. L.A.S.A: analysis and writing - review & editing. G.G.C.: supervision and writing - review & editing. M.R.C.R.L.: supervision and writing - review & editing. D.B.P.: conceptualization, funding acquisition, project administration, writing - original draft.

## Supporting Information

Supplementary material: DNA sequences, plasmids, and strains used in this work.

Supplementary data: Data 1 - Raw and processed data for P_*veg*_ and derived promoters test; Data 2 - Raw and processed data for complete circuit test; Data 3 - Raw and processed data for circuit test changing the carbon source; Data 4 - Raw and processed data for scale-up simulation test.

## Acknowledgment

This work was supported by the São Paulo Research Foundation (FAPESP) [grants 2014/17564-7, 2018/04703-0, and 2020/16058-1]; Conselho Nacional de Desenvolvimento Científico e Tecnológico (CNPq) [grants 141389/2021-4, 405490/2021-6, 310023/2020-3, and INCT BioSyn]; and Coordenação de Aperfeiçoamento de Pessoal de Nível Superior - Brasil (CAPES) [Finance Code 001].

