## Supplementary tables and figure for "Preventing production escape during scale-up using an engineered glucose-inducible genetic circuit"

**Supplementary Table 1**. Promoter sequences engineered in this work.

| **Name** | **Sequence** |
| --- | --- |
| **P*_veg-tetO_*** | *AATTTTGTCAAAATAATTTTATTGACAACGTCTTATTAACGTTGATACCGGTTAAATTTTATTTGACAAAAATGGGCTCGTGTTGTACAATAAATGT*ACTCTATCATTGATAGAG**T**TCCGGCAAAAAAGGGCAAGGTGTCCTAGTAAGGTCGAC**AGGAGGA**TACTAG |
| **P*_veg-cre_*** | *AATTTTGTCAAAATAATTTTATTGACAACGTCTTATTAACGTTGATACCGGTTAAATTTTATTTGACAAAAATGGGCTCGTGTTGTACAATAAATGT*ATTTTGTTAACGCTTACAATTTTGTTAACGCTTACATTTATACTAGAGAA**AGAGGAGA**AATACTAG |

P*_veg_* is in italics, *tetO* site and double *cre* sites are gray shaded, and the RBS is bold.

**Supplementary Table 2**. List of plasmids used.

| **Plasmid** | **Host** | **Description** | **Source** |
| --- | --- | --- | --- |
| pBS3C*lux* | *E. coli* Top10 | Plasmid carrying the promoterless *luxABCDE* cluster integrates at the *sacA* locus in the *B. subtilis* genome | BGSC ECE261 |
| pBS4S | *E. coli* Top10 | Empty plasmid, integrates at the *thrC* locus in *B. subtilis* | BGSC 259 |
| pSB1C3-TetR-LVA | *E. coli* Top10 | Plasmid carrying the tetR fused to the LVA degradation tag | BBa_P0440 |
| pSB1C3-T7RNAP | *E. coli* Top10 | T7 RNA polymerase | BBa_K145001 |
| pSB1C3-P*_veg_*_-cre_-TetR | *E. coli* Top10 | tetR-LVA under the control of the P*_veg_*_-cre_ | This work |
| pBS3C-P*_veg_*-*lux* | *E. coli* Top10 | *luxABCDE* under the control of the P*_veg_* | This work |
| pBS3C-P_T7_-*lux* | *E. coli* Top10 | *luxABCDE* under the control of the P_T7_ | This work |
| pBS3C-P*_veg_*_-_*_tet_*_O_-*lux* | *E. coli* Top10 | *luxABCDE* under the control of the P*_veg_*_-_*_tet_*_O_ | This work |
| pBS3C-P*_veg_*_-cre_-*lux* | *E. coli* Top10 | *luxABCDE* under the control of the P*_veg_*_-cre_ | This work |
| pBS4S-P*_veg_*_-_*_tet_*_O_-T7RNAp | *E. coli* JM109 | T7 RNA polymerase under the control of the P*_veg_*_-_*_tet_*_O_ | This work |
| pBS4S-P*_veg_*_-_*_tet_*_O_-T7RNAp–P*_veg_*_-_*_cre_*-TetR | *E. coli* JM109 | Glucose-inducible circuit | This work |

**Supplementary Table 3**. List of strains used.

| **Strain** | **Description** | **Source** |
| --- | --- | --- |
| *E. coli* Top10 | Cloning strain | Lab stock |
| *E. coli* JM109 | Cloning strain | Lab stock |
| *E. coli* BL21(DE3) | T7 RNA polymerase-driven expression | Lab stock |
| *B. subtilis* 168 | Wild-type, *trpC2* | Lab stock |
| *E. coli* pBS3C-P_T7_-*lux* | *E. coli* BL21 expressing the lux operon from the T7 promoter | This work |
| *B. subtilis* :: *lux* | *B. subtilis* 168 *sacA* :: pBS3C*lux* | This work |
| *B. subtilis* :: P_T7_-*lux* | *B. subtilis* 168 *sacA* :: pBS3C-P_T7_-*lux* | This work |
| *B. subtilis* :: P*_veg_*-*lux* | *B. subtilis* 168 *sacA* :: pBS3C-P*_veg_*-*lux* | This work |
| *B. subtilis* :: P*_veg_*_-tetO_-*lux* | *B. subtilis* 168 *sacA* :: pBS3C-P*_veg-tetO_*-*lux* | This work |
| *B. subtilis* :: P*_veg_*_-cre_-*lux* | *B. subtilis* 168 *sacA* :: pBS3C-P*_veg-cre_*-*lux* | This work |
| *B. subtilis* :: P_T7_-*lux ::* P*_veg_*_-_*_tet_*_O_-T7RNAp | *B. subtilis* :: P_T7_-*lux thrC ::* pBS4S-P*_veg_*_-_*_tet_*_O_-T7RNAp | This work |
| *B. subtilis* *::* P*_veg_*_-_*_tet_*_O_-T7RNAp–P*_veg_*_-_*_cre_*-TetR | *B. subtilis* 168 *thrC ::* pBS4S-P*_veg_*_-_*_tet_*_O_-T7RNAp–P*_veg_*_-_*_cre_*-TetR | This work |
| *B. subtilis*::P_T7_-*lux* :: P*_veg_*_-_*_tet_*_O_-T7RNAp–P*_veg_*_-_*_cre_*-TetR | *B. subtilis*::P_T7_-*lux thrC ::* pBS4S-P*_veg_*_-_*_tet_*_O_-T7RNAp–P*_veg_*_-_*_cre_*-TetR | This work |

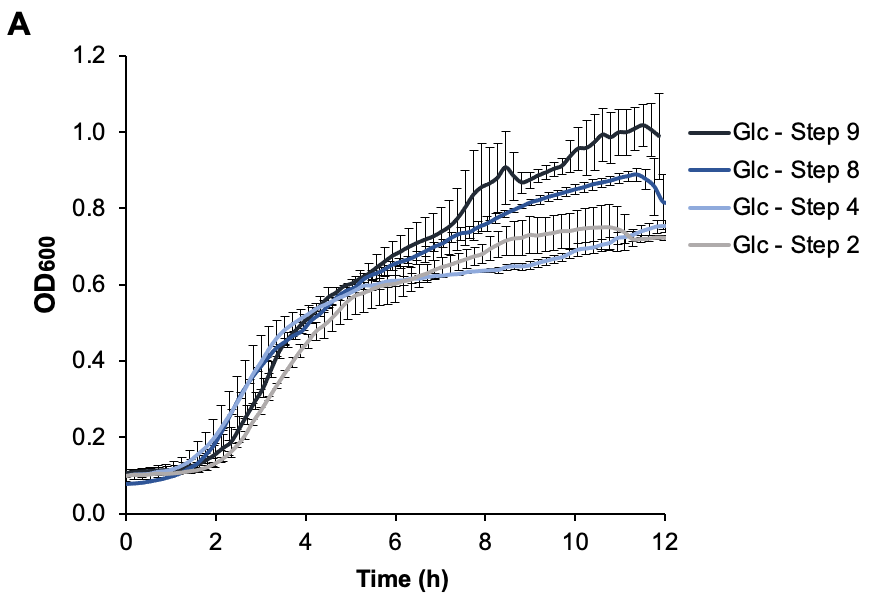

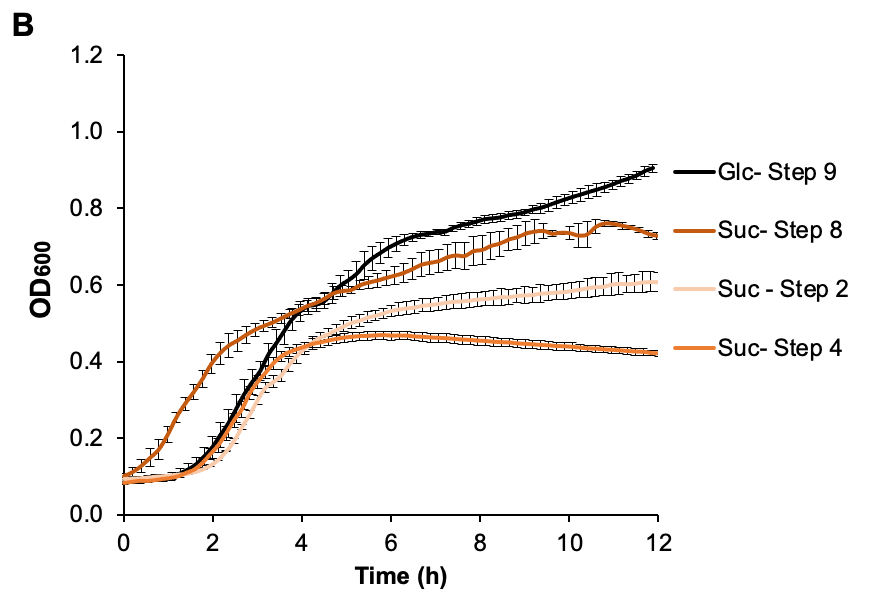

**Supplementary Figure 1**. Growth curves measured during the simulated production scale-up through nine serial cultivation transfers (≥ 55 generations). Two treatments were applied, continuous glucose feeding for nine steps (A) and eight steps of sucrose feeding followed by a glucose feeding ninth step (B). Growth curves are presented for steps 2, 4, 8, and 9 (see Fig. 3B for more information). The ninth step simulates the final large-scale production cultivation. Results are presented as the mean of biological triplicates and the standard error.
